# An *in vitro* model of fibrosis using crosslinked native extracellular matrix-derived hydrogels to modulate biomechanics without changing composition

**DOI:** 10.1101/2022.02.02.478812

**Authors:** M. Nizamoglu, R.H.J de Hilster, F. Zhao, P.K. Sharma, T. Borghuis, M.C. Harmsen, J.K. Burgess

## Abstract

Extracellular matrix (ECM) is a dynamic network of proteins, proteoglycans and glycosaminoglycans, providing structure to the tissue and biochemical and biomechanical instructions to the resident cells. In fibrosis, the composition and the organization of the ECM are altered, and these changes influence cellular behaviour. Biochemical (i. e. protein composition) and biomechanical changes in ECM take place simultaneously *in vivo*. Investigating these changes individually *in vitro* to examine their (patho)physiological effects has been difficult. In this study, we generated an *in vitro* model to reflect the altered mechanics of a fibrotic microenvironment through applying fibre crosslinking via ruthenium/sodium persulfate crosslinking on native lung ECM-derived hydrogels. Crosslinking of the hydrogels without changing the biochemical composition of the ECM resulted in increased stiffness and decreased viscoelastic stress relaxation. The altered stress relaxation behaviour was explained using a generalized Maxwell model. Fibre analysis of the hydrogels showed that crosslinked hydrogels had a higher percentage of matrix with a high density and a shorter average fibre length. Fibroblasts seeded on ruthenium-crosslinked lung ECM-derived hydrogels showed myofibroblastic differentiation with a loss of spindle-like morphology together with greater α-smooth muscle actin (α-SMA) expression, increased nuclear area and circularity without any decrease in the viability, compared with the fibroblasts seeded on the native lung-derived ECM hydrogels. In summary, ruthenium crosslinking of native ECM-derived hydrogels provides an exciting opportunity to alter the biomechanical properties of the ECM-derived hydrogels while maintaining the protein composition of the ECM to study the influence of mechanics during fibrotic lung diseases.

## Introduction

Extracellular matrix (ECM) is the structural component of every tissue, formed by a complex network of proteins, glycosaminoglycans and proteoglycans. ^1^ The highly tissue-specific nature of the ECM is dictated by the presence of a defined grouping of matrisome elements, incorporating demarcated ratios of ECM proteins. ^2^ These distinctions also result in different mechanical properties of the ECM, depending on the origin of the tissue. ^3^ Next to being structural support for the cells, the ECM provides biochemical and biomechanical cues to cells *in vivo*. ^4^ As such, it has proven challenging to mimic and incorporate the ECM structure and mechanics into (*in vitro)* studies regarding the structure and function of the ECM in health and disease. In fibrotic diseases, not only is the ECM composition altered but also its mechanical properties, resulting in higher stiffness and decreases in stress relaxation. ^5^ All the changes that are evident within a fibrotic ECM have been revealed to instruct cells and influence their responses to contribute to the progression of fibrosis, as reported and reviewed elsewhere. ^6–12^ To investigate the mechanical properties of (fibrotic) ECM *in vitro*, the ECM is often mimicked using hydrogels. ECM-derived hydrogels, which have been introduced to the field in the last decade, are a promising alternative to other types of hydrogels such as collagen, gelatine, or hyaluronic acid. ^13^ ECM hydrogels which are developed from native decellularized tissue, retain most of the native ECM composition and, in general, resemble the mechanical properties of the parent tissue. ^14^ The most common method to produce hydrogels from ECM is to digest decellularized ECM powder with porcine pepsin at low pH with constant agitation. ^13^ Our recent study illustrated the preparation of ECM-derived hydrogels from human decellularized lung ECM, and established that the mechanical properties of the diseased (fibrotic) lung ECM-derived hydrogels resembled the mechanics of the decellularized fibrotic lung ECM. ^14^ Fibrotic lung ECM (both in native and hydrogel form) showed decreased viscoelastic stress relaxation compared to control lung ECM. ^14^ The stiffness of fibrotic lung tissue was ~10 times higher than its hydrogel counterpart, possibly due to the absence of chemical crosslinks and lung-resident cells in the ECM hydrogel. Previous studies showed that the composition of fibrotic lung ECM is different to that seen in control lung due to dysregulation of the ECM degradation/deposition processes resulting in an aberrant ECM. ^15^ To investigate the separate influences on the cells of the altered mechanical properties or ECM composition in the fibrotic microenvironment, novel *in vitro* models are needed. Recently, altering the mechanical properties of methacrylate or thiol functionalized ECM-derived hydrogels using click-chemistry has been shown. ^16, 17^ Given that these processes rely on interactions with amine groups of lysine or arginine amino acids, which are known to be parts of cell binding domains including GFOGER, IKVAV or RGD. The implications of methacrylation or thiolation of ECM proteins on cellular functions still need to be explored. ^16, 18–20^ Alternatively, chemical crosslinking has been applied to ECM-derived hydrogels using harsh chemicals such as glutaraldehyde or genipin, but cytotoxicity limits their use when cells are present in the hydrogel. ^21^ Another option is using near visible light UV-induced ruthenium/sodium persulfate (SPS) crosslinking, which has been employed on several other types of hydrogels (gelatine or fibrin) with and without cells present in the hydrogels. ^22, 23^ The higher wavelength (405 nm) of the crosslinking light, which decreases the cytotoxicity, and the lack of requirement for any additional functionalization on the target material are the main advantages of this crosslinking method. ^24^ Using ruthenium/SPS crosslinking to reinforce the mechanical stability of the ECM-derived hydrogels has recently been reported by Kim et al. ^25^; however, the implications of altering the mechanical properties of the hydrogels without changing the (bio)chemical composition have yet to be explored in terms of fibrosis and for developing *in vitro* models for fibrosis research.

In this study, we aimed to develop an *in vitro* model for examining the influence of mechanical properties of the fibrotic microenvironment by using native lung-derived ECM hydrogels (LdECM), which were generated using ruthenium/SPS crosslinking. We hypothesized that the ruthenium crosslinking would increase the stiffness of lung-derived ECM hydrogels (Ru-LdECM), while the viscoelastic relaxation would decrease, to then trigger pro-fibrotic activation of lung fibroblasts.

## Materials and methods

### Porcine Lung Decellularization

Porcine lungs (~6-month, female) were purchased from a local slaughterhouse (Kroon Vlees, Groningen, the Netherlands). The lung was dissected, cartilaginous airways and large blood vessels removed, before cutting into ~1cm^3^ cubes that were homogenized in a kitchen blender prior to decellularization. The lung homogenate was decellularized as previously described. ^26, 27^ In short, the homogenate was repeatedly washed with Milli-Q^®^ water and centrifuged at 3,000 x g until the supernatant was completely clear. The sedimented material went through two rounds of sequential treatment with 0.1 % Triton X-100 (Sigma-Aldrich, St. Louis, MO, USA), 2 % sodium deoxycholate (Sigma-Aldrich), 1 M NaCl solution and 30 μg/mL DNase (Sigma-Aldrich) in MgSO_4_ (Sigma-Aldrich) 1.3 mM and CaCl_2_ (Sigma-Aldrich) 2 mM, 10 mM Tris pH8 (Sigma-Aldrich) solution each for 24 h at 4°C with constant shaking, except for the DNAse treatments, which were at 37°C with shaking. The volume ratio of tissue homogenate to decellularization/washing solution was always 1:10. Between treatments, the homogenate was washed three times with Milli-Q^®^ water, with centrifugation at 3,000 x g between washes. After two cycles of decellularization, the tissue homogenate was sterilised by adding 0.18 % peracetic acid and 4.8 % ethanol, and left shaking at 4 °C for 24 h. After tissue sterilization the resultant decellularized ECM was washed three times with sterile Dulbecco’s phosphate-buffered saline (DPBS) and stored in sterile DPBS containing 1 % penicillin-streptomycin (Gibco Invitrogen, Carlsbad, CA, USA) at 4 °C (**Figure 1A**).

**Figure 1:**
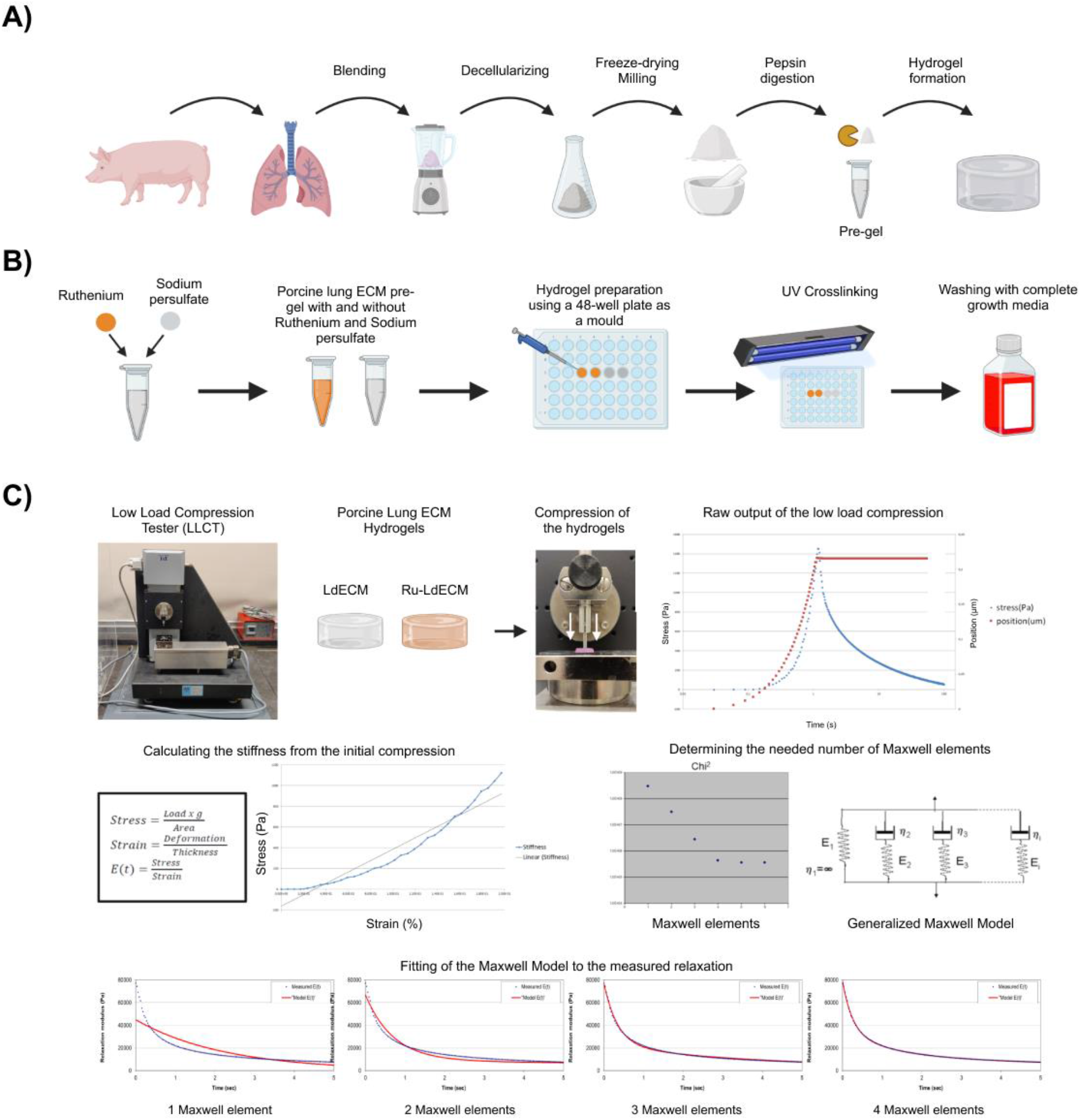
Schematic representation of the methodology. A) Porcine lung ECM hydrogel preparation. Porcine lungs were blended, decellularized and freeze-dried before grinding to a fine powder. Afterwards, the ECM powder was pepsin digested to prepare the pre-gel solution which can form hydrogels after incubating at 37°C. B) Fibre crosslinking of porcine lung ECM hydrogels. Lung derived-ECM (LdECM) hydrogels were used as is or mixed with ruthenium and sodium persulfate solutions before casting to 48-well plates. Afterwards, the pre-gel solutions were incubated at 37°C and UV-crosslinked. C) Mechanical characterization of the uncrosslinked and ruthenium crosslinked ECM hydrogels. Low Load Compression Tester (LLCT) was used to determine the stiffness and stress relaxation of LdECM and Ru-LdECM hydrogels. The stress relaxation was modelled with a Maxwell elements system, and a chi^2^ analysis was used to determine the number of Maxwell elements to fit the measured relaxation. Figure created with BioRender.

### Hydrogel preparation

The decellularized lung ECM was snap-frozen in liquid nitrogen and lyophilized with a FreeZone Plus lyophilizer (Labconco, Kansas City, USA), before being ground into a powder with an A11 Analytical mill (IKA, Staufen, Germany). For solubilization, 20 mg/mL of ECM powder was digested with 2 mg/mL porcine pepsin (Sigma-Aldrich) in 0.01 M HCl under constant agitation at RT for 48 h. Digestion was stopped by neutralising the pH with 0.1 M NaOH and the solution was brought to 1X PBS with one-tenth volume 10X PBS to generate the lung ECM pre-gel solution which was stored at 4°C indefinitely.

A ruthenium Visible Light Photo initiator (400-450nm) kit (Advanced BioMatrix, San Diego, California, US) containing pentamethyl cyclopentadienyl bis(triphenylphosphine) ruthenium(II) chloride (CAS Number: 92361-49-4, hereafter referred as ruthenium) and sodium persulfate (CAS: 7775-27-1) was used to crosslink the LdECM hydrogels (**Figure 1B**). 20 μL of each ruthenium. (37.4 mg/mL) and sodium persulfate (119 mg/mL) solutions were added per 1 mL of ECM hydrogel. The control gel received the same volume of sterile ddH_2_O water. In the dark, both the ruthenium-containing gel and the control gel without ruthenium were pipetted (200 μL) into a 48-well plate and incubated at 37°C for 1 h. After the hydrogels had settled, crosslinking of the ECM by photoinitiated ruthenium was triggered by exposing the samples to UV/Visible light from 4.5 cm distance using 2 x 9W UV lamps (405 nm) for 5 min to generate Ru-LdECM hydrogel (**Figure 1B**). Finally, the gels were immersed in 400 μL of Dulbecco’s Modified Eagle Medium (DMEM) Low Glucose growth media (Lonza) supplemented with 10% foetal bovine serum (FBS), 1% penicillin-streptomycin and 1% GlutaMAX (Gibco) (hereafter referred as complete growth medium), and were washed 3X with media before cell seeding in order to remove excess (both reacted and unreacted) ruthenium and sodium persulfate.

### Characterization of the Mechanical properties

Both LdECM and Ru-LdECM hydrogels were made as described in hydrogel preparation. The gels were subjected to uniaxial compression with a 2.5 mm diameter plunger at three different locations, at least 2 mm away from the gel border and ensuring 2 mm or more between each compression site (**Figure 1C**). The stress relaxation test was performed with a low-load compression tester (LLCT) at RT as described previously. ^26, 28, 29^ The LabVIEW 7.1 program was used for the LLCT load cell and linear positioning for control and data acquisition. The resolution in position, load, and time determination was 0.001 mm, 2 mg, and 25 ms, respectively, and the compression speed was controlled in feedback mode. Samples were compressed to 20% of their original thickness (strain ε = 0.2) at a deformation speed of 20 %/s (strain rate 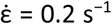). The deformation was held constant for 100 s and the stress continuously monitored. During compression, the required stress was plotted against the strain. In this plot, a linear increase in stress as a function of strain was observed between a strain of 0.04 and 0.1; the slope of the line fit to this region was taken as stiffness (Young’s modulus). Since the stiffness of the viscoelastic gel depends on the strain rate, values reported here are valid only at a strain rate of 0.2 s^−1^.

After compression, the required stress to maintain a constant strain of 0.2 s^−1^, continuously decreases with time, which is a clear indication of the viscoelastic nature of the hydrogels and called stress relaxation. The shape of the stress relaxation curve was mathematically modelled with a generalized Maxwell model (2) (Fig. 1C). The continuously changing stress [*σ(t)*] was converted into continuously changing stiffness [*E(t)*] by dividing with the constant strain of 0.2 s^−1^. Obtained *E(t)* values were fitted to Eq. 1 to obtain the relaxation time constants (*τ_i_*), and Eq. 2 provided relative importance (*R_i_*) for each Maxwell element.

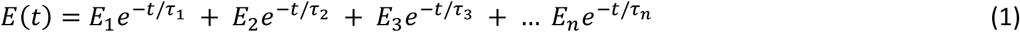

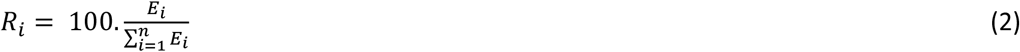

where *i* varies from 1 to 4 or from 1 to 3 when necessary. The optimal number of Maxwell elements was determined with the chi-square function expressed by Eq. 3 (typically 3 or 4) and visually matching the modelled stress relaxation curve to the measured curve (Fig. 1C).

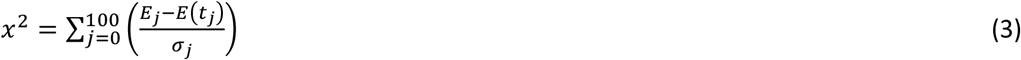

where *j* varies from 0 to 100 s, *E_j_* is the experimentally measured value at time *j*, *E(t_j_)* is the fit value at time *j* calculated with Eq. 1, and *σ_j_* is the standard error that the LLCT makes because of inherent errors in position, time, and load measurements.

### Histological characterisation of ECM hydrogel fibre structure

LdECM and Ru-LdECM hydrogels were prepared and washed as described above and fixed with 2% paraformaldehyde in PBS (PFA; Sigma-Aldrich) at RT for 20 min. The gels were then embedded in 1% Ultrapure agarose (Invitrogen, Waltham, MA, USA) before using a graded alcohol series to dehydrate followed by paraffin embedding. Sections (4 μm) were deparaffinised, and stained with 0.1 % Picrosirius Red (PSR) (Sigma-Aldrich) in 1.3% aqueous solution of picric acid to visualize collagens and their network. Slides were mounted with Neo-Mount^®^ Mounting Medium (Merck, Darmstadt, Germany).

### Cell culture

MRC-5 foetal lung fibroblasts (n=5) were cultured in complete growth medium. The MRC-5s were washed with Hank’s Balanced Salt Solution (HBSS; Gibco), harvested using 0.25% Trypsin-EDTA (Gibco) and centrifuged at 500 x g for 5 minutes. Cells were resuspended in 1 mL complete growth media and counted with a NucleoCounter NC-200™ (Chemometec, Allerod, Denmark). Fibroblasts were seeded on top of pre-prepared and washed LdECM and Ru-LdECM hydrogels in complete growth media with the seeding density 10.000 cells/gel. The cells were cultured on the gels for 1 or 7 days. Gels used for live/dead staining were stained with the live/dead stain and were subsequently harvested. Gels intended for immunofluorescent imaging were fixed in 2% PFA in PBS for 30 minutes. After fixation, hydrogels were washed three times with PBS and stored in PBS containing 1% penicillin-streptomycin at 4°C until analyses.

### Live/dead staining

Cell viability of the MRC-5 cells cultured on LdECM and Ru-LdECM hydrogels was assessed after 1 and 7 days using Calcein AM (Thermo Scientific, Breda, the Netherlands) to stain live cells and propidium iodide (PI; Sigma-Aldrich) for staining dead cells, as previously described. ^30^ The hydrogels were first washed with HBSS and then incubated with serum free media containing 5 μM Calcein AM and 2 μM PI, for 1 hour at 37°C. After incubation, fluorescent images were captured using a EVOS Cell Imaging System (Thermo Scientific) with GFP (509 nm) and Texas Red (615 nm) channels.

### Immunofluorescence Staining

The hydrogels were treated with avidin/biotin blocking kit (ThermoFisher) before being incubated with 0.5 μg/mL biotinylated wheat germ agglutinin (Vector Laboratories, Burlingame, USA) for 20 min at 37°C. Then the hydrogels were washed and permeabilized by incubating with 0.5% v/v Triton X-100 in HBSS for 10 min at RT and subsequently blocked in 2.5% v/v BSA + 0.1% Triton-X 100 in HBSS for 30 min at RT. Endogenous peroxidase activity was blocked by 30 min incubation in a 0.3% hydrogen peroxide solution. Afterwards, the hydrogels were incubated overnight with a mouse anti-human α-smooth muscle actin antibody (DAKO, Glostrup, Denmark) at 4°C. A rabbit-anti-mouse antibody conjugated with peroxidase (DAKO) and streptavidin conjugated with Alexa Fluor 555 (ThermoFisher) were used as a second step for 45 minutes at room temperature Staining for α-SMA was then developed by Opal650 tyramide (Akoya Biosciences, Marlborough MA, USA) according to the manufacturer’s instructions. After staining with 0.1 μg/mL DAPI solution (Merck), the hydrogels were mounted with Citifluor Mounting Medium (Science Services, Munich, Germany) and fluorescence microscopy was performed to acquire images.

### Imaging and image analysis

Fluorescent images of PSR-stained LdECM and Ru-LdECM hydrogel sections were generated with Zeiss LSM 780 CLSM confocal microscope (Carl Zeiss NTS GmbH, Oberkochen, Germany), λ_ex_ 561 nm / λ_em_ 566/670 nm at 40x magnification. TWOMBLI plugin for FIJI ImageJ was used to assess the number of fibres, end points, branching points, total fibre length and alignment, lacunarity, high density matrix (HDM) and curvature of the fibres as previously described (*Supplementary Figure 1*). ^26, 31^ Fluorescent images of cell-seeded hydrogels stained for αSMA, wheat germ agglutinin and DAPI were generated with Leica SP8 confocal microscope (Leica, Wetzlar, Germany), using λ_ex_ 627 nm / λ_em_ 650 nm for αSMA, λ_ex_ 555 nm / λ_em_ 580 nm for wheat germ agglutinin and λ_ex_ 359 nm / λ_em_ 457 nm for DAPI at 40X and 63X magnifications. CellProfiler 4.2.1 software was used to analyse the nuclei characteristics on the DAPI-stained images as previously described. ^32^ Five separate images per sample (n = 5) were used to calculate the stiffness-induced changes in the nuclei area and eccentricity (which is also described as inverse circularity) (Supplementary Figure 2). Circularity of the samples were calculated from the eccentricity values using the equation (4).

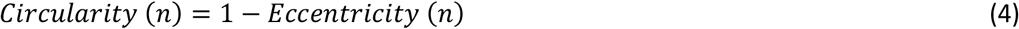

### Statistical analysis

All statistical analyses were performed using GraphPad Prism v9.1.0 (GraphPad Company, San Diego, USA). Data are presented as mean values with standard deviation (SD). All data was tested for outliers using the robust regression and outlier removal (ROUT) test and analysed for normality using Shapiro-Wilk and Q-Q plots (*Supplementary Figure 3*). For the data that were normally distributed, differences between control porcine lung ECM hydrogel and ruthenium crosslinked ECM hydrogels were tested by paired t-test to compare the effect of crosslinking between different experiments. For the data that were not normally distributed, Mann-Whitney test was used to compare the effect of crosslinking between different experiments. All data were considered significantly different when p < 0.05.

## Results

### Ruthenium crosslinking increases hydrogel stiffness

Both LdECM and Ru-LdECM solutions were able to form hydrogels after incubating at 37 °C. UV/Visible light crosslinking did not result in a macroscopic change in the LdECM hydrogels. Ru-LdECM hydrogels (and the solution before the crosslinking) had a bright orange colour due to the ruthenium addition. Stiffness measurements on LdECM and Ru-LdECM hydrogels were performed using a low-load compression tester (LLCT). Ruthenium crosslinking increased the stiffness of the Ru-LdECM 5-10-fold (p = 0.0026, paired t-test) (**Figure 2**).

**Figure 2:**
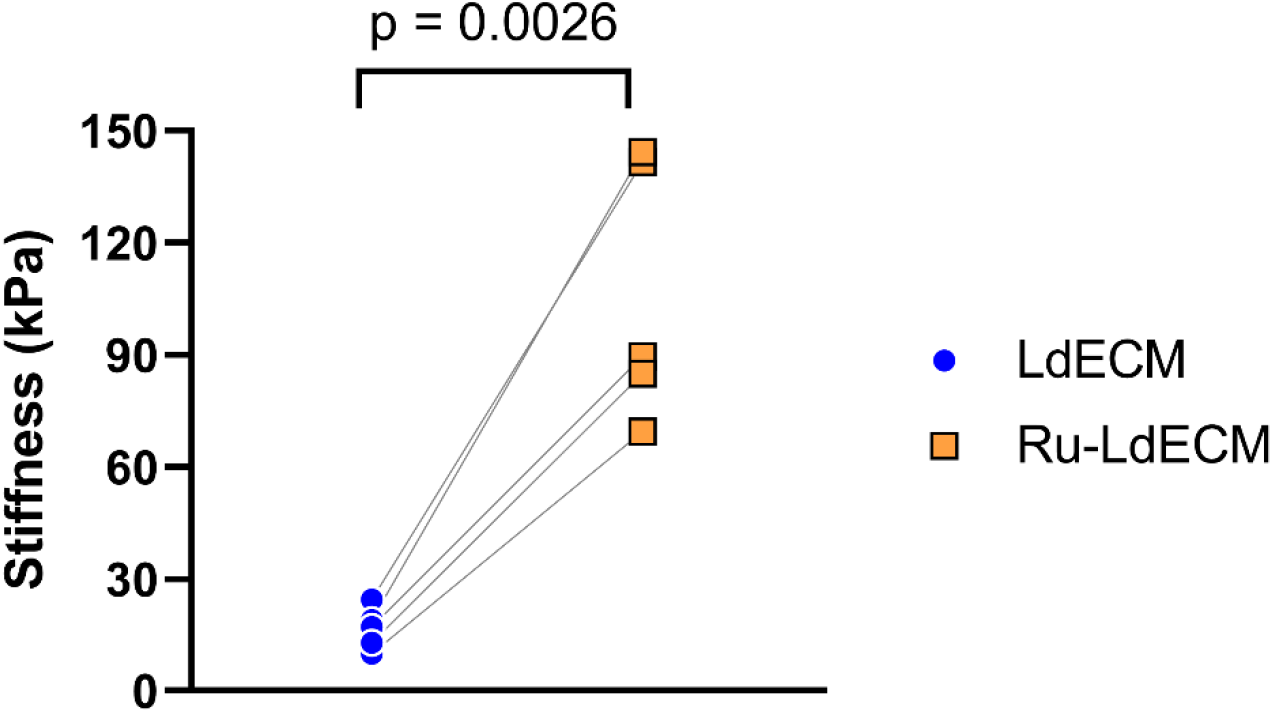
Comparison of stiffness of control and ruthenium-crosslinked hydrogels. LdECM and Ru-LdECM hydrogels were mechanically tested using Low Load Compression Tester (LLCT) with a fixed 20% strain ratio. Each dot represents the mean of three independent measurements on the same hydrogel for each sample (n = 5). Applied test: Paired t-test. LdECM: Lung-derived ECM Hydrogels, Ru-LdECM: Ruthenium-crosslinked Lung-derived ECM Hydrogels

### Decreased stress relaxation rate in ruthenium-crosslinked ECM hydrogels

The stress relaxation behaviour of both the LdECM and RU-LdECM hydrogels were measured after applying 20% strain using LLCT measurement. The average stress relaxation profiles of both groups over 100 s are visualized in **Figure 3A**. Ru-LdECM hydrogels did not reach 100% stress relaxation during the 100 s monitored, while some LdECM hydrogels achieved 100% stress relaxation. In addition to the decreased total stress relaxation percentage (in 100s) in the Ru-LdECM hydrogels, the relaxation profile was different. The rate of stress relaxation slowed down earlier in the crosslinked hydrogels. To assess the dynamic differences in the initial stress relaxation behaviour patterns in both groups the time to reach 50% total stress relaxation was compared. LdECM hydrogels reached 50% stress relaxation in significantly shorter time compared to the Ru-LdECM hydrogels. (p = 0.0054, paired t-test) (**Figure 3B**).

**Figure 3:**
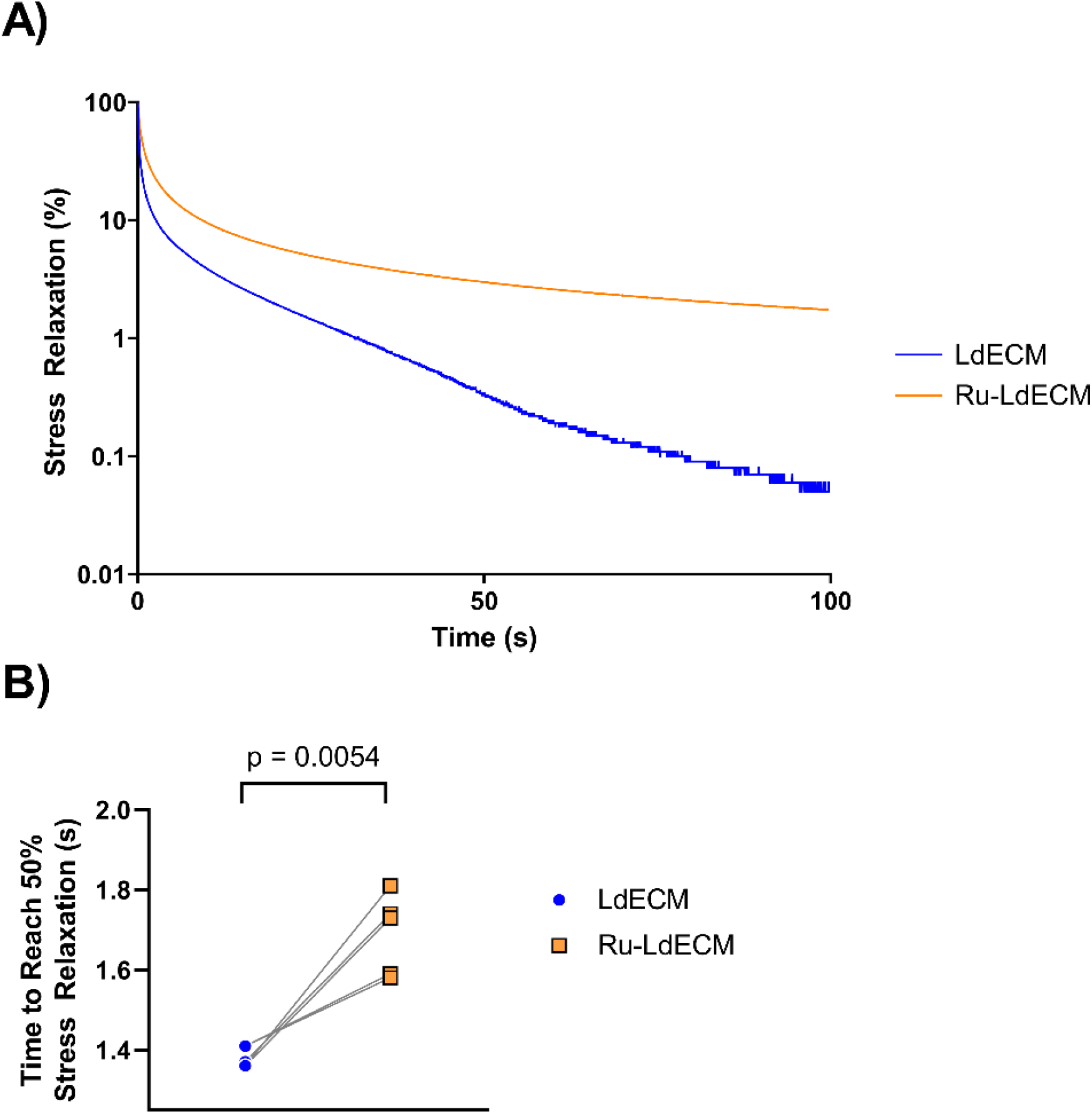
Stress relaxation in control and ruthenium-crosslinked ECM-derived hydrogels. After compressing the LdECM and Ru-LdECM hydrogels using Low Load Compression Tester (LLCT) with a fixed 20% strain ratio, the stress relaxation behaviour was recorded over 100s period. A) Average stress relaxation behaviour over 100 s duration. B) Time taken to reach 50% stress relaxation. Each dot represents the mean of three independent measurements on the same hydrogel for each sample (n = 5). Applied test: Paired t-test. LdECM: Lung-derived Extracellular Matrix Hydrogels, Ru-LdECM: Ruthenium-crosslinked Lung-derived Extracellular Matrix Hydrogels.

### Altered relaxation profile in ruthenium-crosslinked ECM hydrogels

Since the ECM hydrogels are a viscoelastic material with various elastic (e.g., ECM proteins) and viscous components (e.g., water, bound water), we mathematically modelled this using a generalized Maxwell model. This approach allowed the total relaxation data to be split into Maxwell elements that can theoretically be attributed to physical components in the hydrogels. Each of these Maxwell elements are responsible for a part of the total relaxation (relative importance), as well as occurring within a specific time window during the relaxation process. The distribution of the time constants of these different elements and their respective relative importance are presented in **Figure 4**. The stress relaxation of the LdECM hydrogels could be modelled with 3 Maxwell elements while Ru-LdECM hydrogels needed 4 Maxwell elements. Next to the difference in the number of Maxwell elements required to explain the relaxation profiles, the time constants of the elements differed between the two groups: the first element was significantly faster in the LdECM hydrogels (p = 0.002, paired t-test) while the third element took longer in the LdECM hydrogels (p = 0.002, paired t-test) compared to the Ru-LdECM (**Figure 4A**).

**Figure 4:**
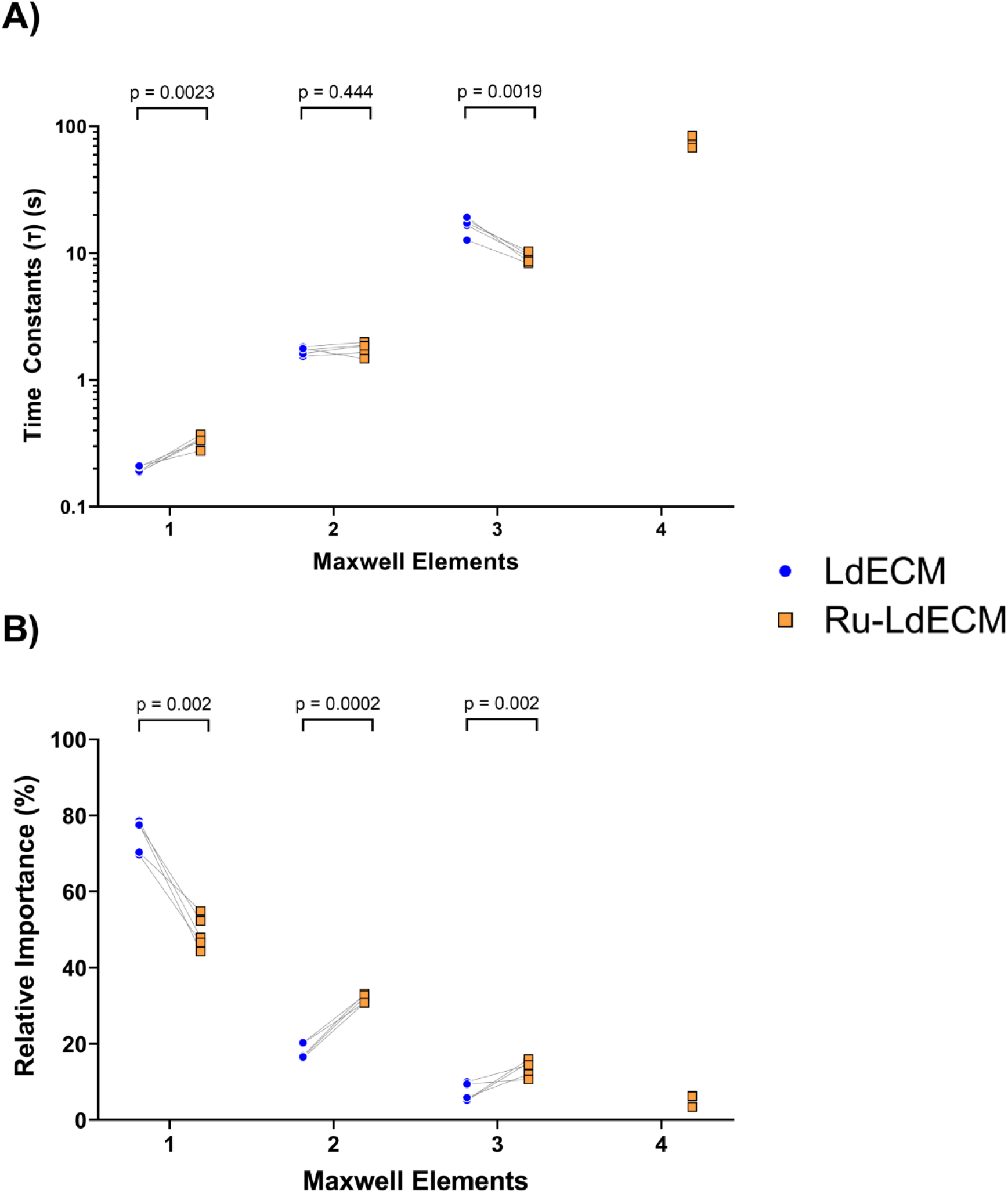
Analysis of the stress relaxation behaviour through the generalized Maxwell model system. The relaxation profiles of the both types of hydrogels over 100 s period was mathematically modelled using a Maxwell model system and the relative importance values of the Maxwell Elements were determined. A) Time constants for each Maxwell element for LdECM and Ru-LdECM hydrogels. B) Relative importance (%) of the each Maxwell element for LdECM and Ru-LdECM hydrogels. Each dot represents the mean of three independent measurements on the same hydrogel (n = 5). Applied test: Paired t-test. LdECM: Lung-derived ECM Hydrogels, Ru-LdECM: Ruthenium-crosslinked Lung-derived ECM Hydrogels

The relative importance of the Maxwell elements was used to assess the individual contribution of each element to the total stress relaxation over the 100 seconds (**Figure 4B)**. In both types of hydrogels, the first element made the greatest contribution to the stress relaxation profile, although the percentage contribution was significantly lower in the Ru-LdECM hydrogels compared with the LdECM hydrogels (p = 0.002, paired t-test). The contribution of the second Maxwell element was the second largest in both groups while the relaxation profile of the Ru-LdECM hydrogels had a significant increase in the contribution of this element compared to uncrosslinked hydrogels (p = 0.0002, paired t-test). The third element had the lowest percentage contribution in LdECM hydrogels, with this contribution being lower than this element in its ruthenium-crosslinked counterpart (p = 0.002, paired t-test).

### Increased density and decreased alignment of fibres in lung ECM hydrogels after ruthenium crosslinking

Ru-LdECM hydrogels had a denser fibre network compared to LdECM hydrogels (**Figure 5A**). The average fibre length was shorter in Ru-LdECM than LdECM hydrogels (p = 0.0026, paired t-test). While the normalized numbers of endpoints and branchpoints did not differ between LdECM and Ru-LdECM hydrogels, the percentage of area with high density matrix (HDM) was greater in Ru-LdECM hydrogels compared with LdECM hydrogels (p = 0.0146, paired t-test). Alignment of the fibres in Ru-LdECM hydrogels was lower than LdECM hydrogels (p = 0.0048, paired t-test) (**Figure 5B-G**).

**Figure 5:**
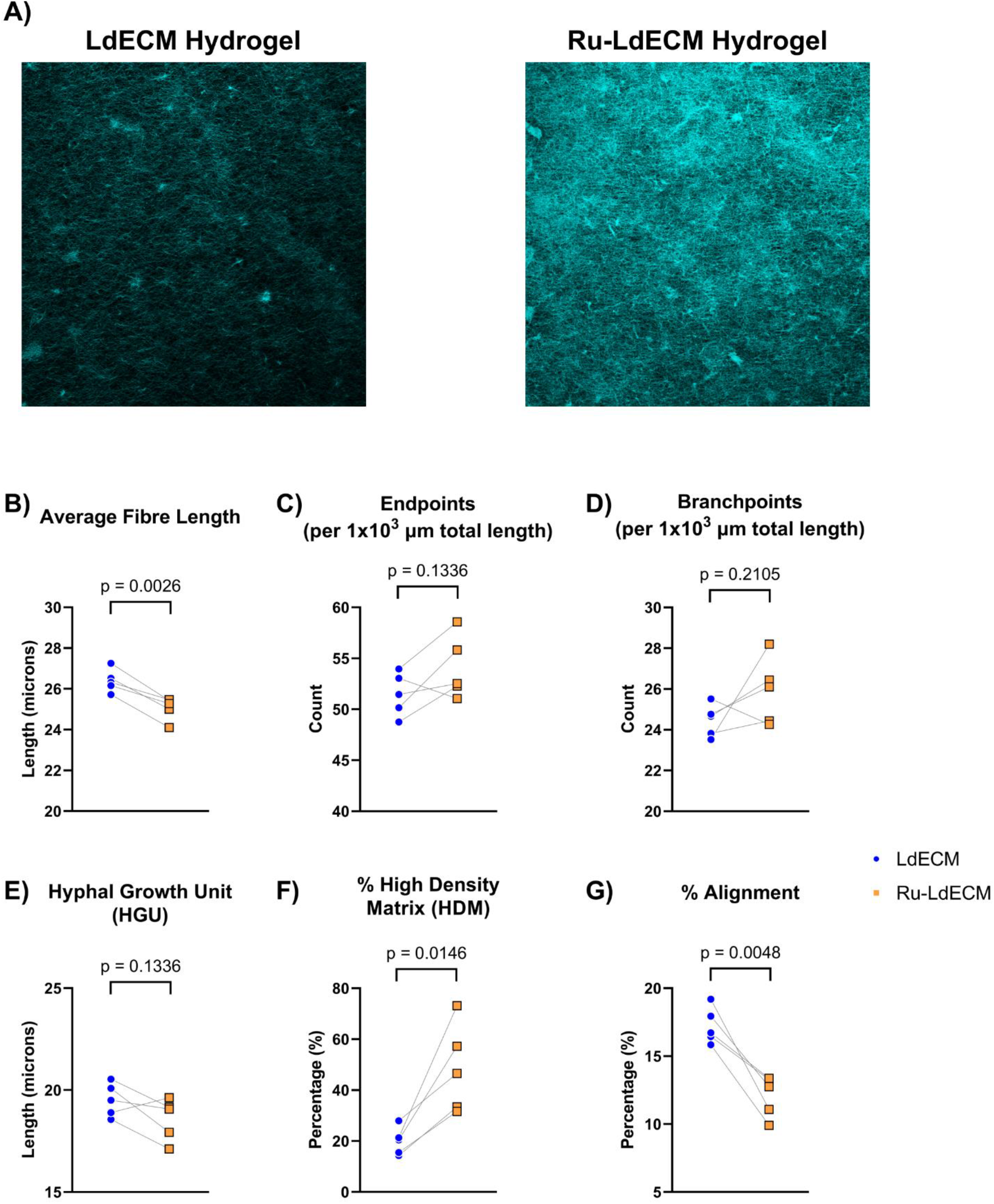
Picrosirius red staining on LdECM and Ru-LdECM and fibre characteristics analysis on the collagen network. LdECM and Ru-LdECM hydrogels were stained using Picrosirius red staining and generated fluorescent images were analysed using TWOMBLI plugin in FIJI ImageJ. A) Representative images of Picrosirius red staining on LdECM and Ru-LdECM hydrogels, B) Average Fibre Length, C) Endpoints per 1000 μm total length, D) Branchpoints per 1000 μm total length, E) HGU, F) % of High Density Matrix (HDM), G) % fibre Alignment. Each dot represents the mean of measurements of 5 different randomized regions on the fluorescent images of Picrosirius red staining for each sample (n=5). Applied statistical test: paired t-test. LdECM: Lung-derived ECM Hydrogels, Ru-LdECM: Ruthenium-crosslinked Lung-derived ECM Hydrogels

The differences in the curvature of the fibres with different length were compared in LdECM and Ru-LdECM hydrogels. Curvature of the fibres with shorter length (< 40 μm) were higher in the crosslinked hydrogels, suggesting that shorter fibres were more bent in Ru-LdECM hydrogels while curvature of the longer fibres was not different between these two groups (**Table 1**).

**Table 1:** TWOMBLI analysis for the curvature of fibres with different lengths. All results show mean ± standard deviation of n = 5. Each analysis was performed using averages of 5 different randomized regions on the fluorescent images of Picrosirius red staining for each sample. LdECM: Lung-derived ECM Hydrogels, Ru-LdECM: Ruthenium-crosslinked Lung-derived ECM Hydrogels. Applied statistical test: paired-t test. P < 0.05 was considered statistically significant.

| Fibre Length | Average Curvature ($\Delta^\circ$) | | P values |
| --- | --- | --- | --- |
|  | LdECM | Ru-LdECM |  |
| 10 $\mu\text{m}$ fibres | 42.40 ± 0.92 | 44.98 ± 0.89 | 0.019 |
| 20 $\mu\text{m}$ fibres | 53.47 ± 0.98 | 56.02 ± 0.92 | 0.022 |
| 30 $\mu\text{m}$ fibres | 59.12 ± 0.71 | 60.84 ± 0.86 | 0.027 |
| 40 $\mu\text{m}$ fibres | 62.55 ± 0.51 | 63.29 ± 1.02 | 0.12 |
| 50 $\mu\text{m}$ fibres | 64.32 ± 0.79 | 64.60 ± 0.84 | 0.58 |

### Ruthenium crosslinking does not affect fibroblast viability but induces altered morphology

After 1 day of culture, no dead cells were observed and the fibroblasts were viable in both types of hydrogels (**Figure 6**). The viability of the fibroblasts did not change over a 7-day culture period (*Supplementary Figure 4*). On both gels the fibroblasts appeared to be lying flat on the surface of the hydrogels; however, the fibroblasts on LdECM hydrogels display a more spindle-shaped morphology, while fibroblasts on Ru-LdECM gels are were more hypertrophic and display more protrusions. Suggesting a more migratory phenotype for fibroblasts on Ru-LdECM hydrogels. At day 7, a fully confluent monolayer was present on both control and crosslinked hydrogels with no differences in viability.

**Figure 6:**
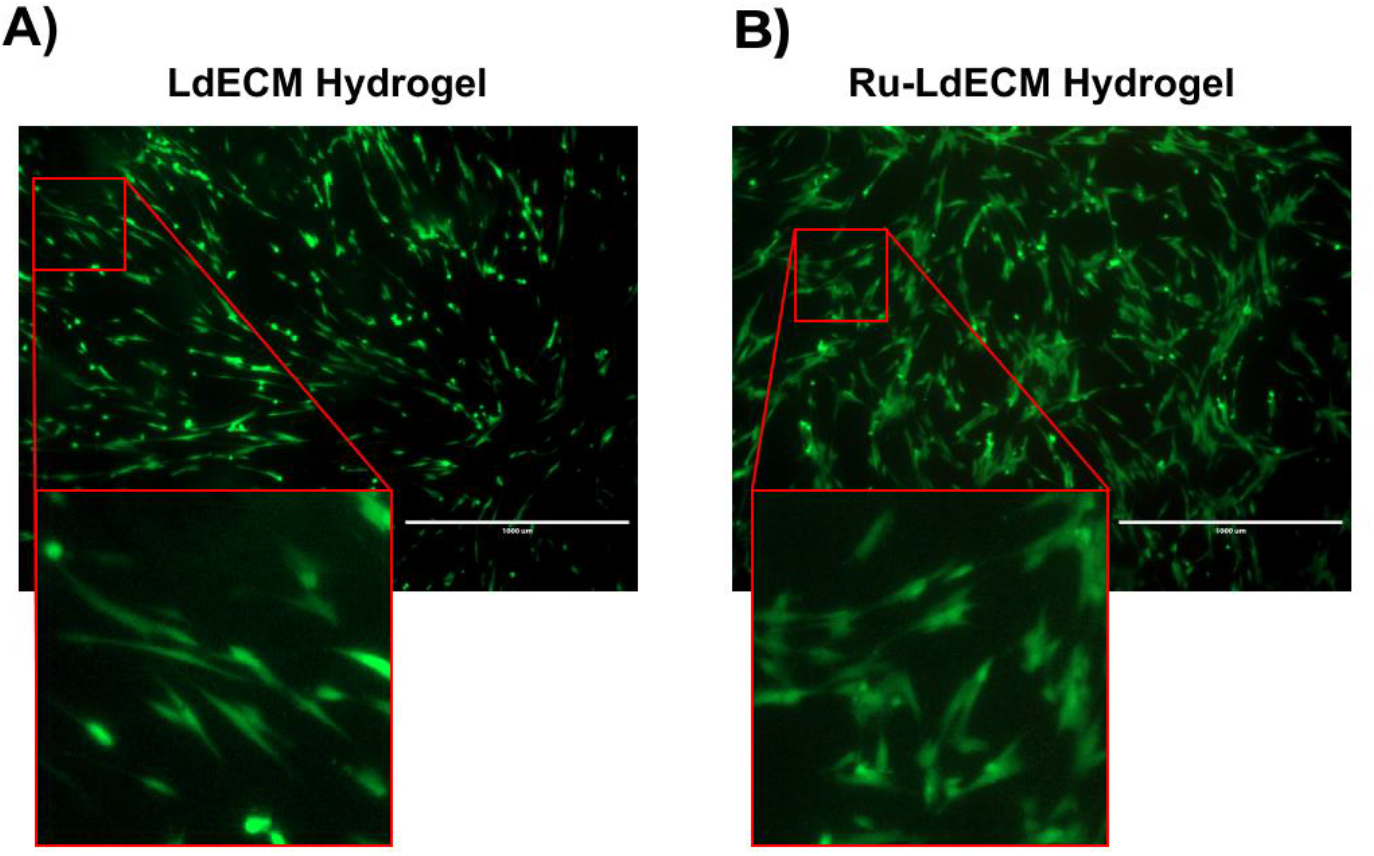
Live/dead staining on MRC-5 fibroblasts seeded on LdECM and Ru-LdECM hydrogels on day 1 using Calcein AM (green) and propidium iodide (red). A) MRC-5 fibroblasts cultured on the LdECM hydrogels, B) MRC-5 fibroblasts cultures on Ru-LdECM hydrogels. Scale bar: 1000 μm. Results are representative for all experiments (n =5). LdECM: Lung-derived ECM Hydrogels, Ru-LdECM: Ruthenium-crosslinked Lung-derived ECM Hydrogels

### Ruthenium crosslinking of ECM hydrogel promotes differentiation of fibroblasts to myofibroblasts

Fibroblasts seeded on Ru-LdECM hydrogels had higher expression of α-SMA when compared to LdECM hydrogel-seeded fibroblasts (**Figure 7**). In addition to the stronger expression of α-SMA, the organization of the cytoskeleton was altered in the fibroblasts seeded on the Ru-LdECM hydrogels (**Figure 7B**, lower row). These myofibroblast-like characteristics were also accompanied by a change in the nuclear morphology.

**Figure 7:**
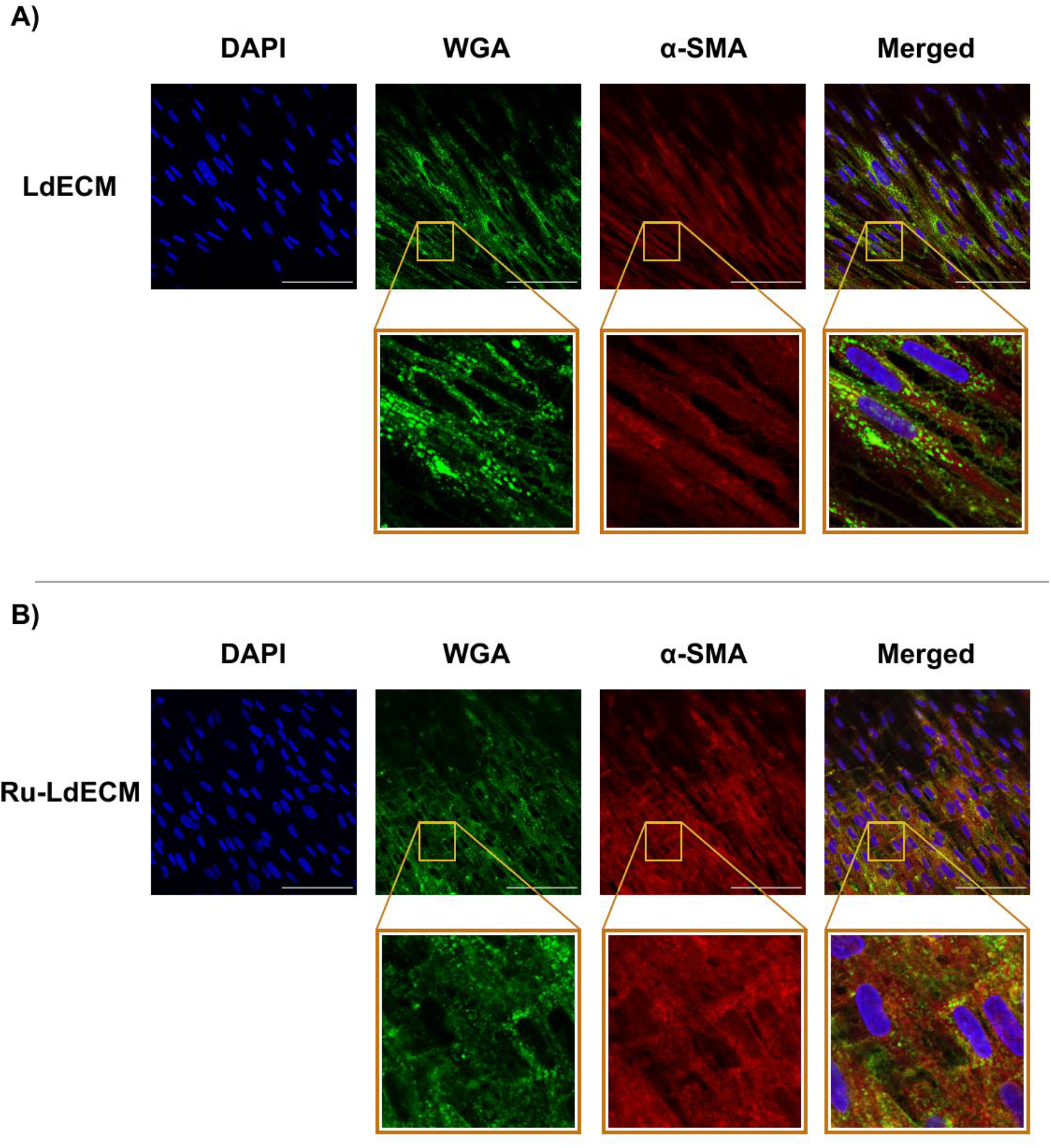
Fluorescence images for the comparison of cell nuclei (DAPI), cell membrane (WGA) and cytoskeleton (a-SMA) on MRC-5 fibroblasts seeded on LdECM and Ru-LdECM hydrogels at day 7. A) Top row: Stained LdECM hydrogels imaged at original objective magnification 40×, bottom row: digitally magnified versions of the respective images B) Stained Ru-LdECM hydrogels imaged at original objective magnification 40×, bottom row: digitally magnified versions of the respective Scale bars: 100 μm. LdECM: Lung-derived ECM Hydrogels, Ru-LdECM: Ruthenium-crosslinked Lung-derived ECM Hydrogels. WGA: Wheat germ agglutinin, α-SMA: alpha smooth muscle actin

At day 7, the nuclei in the fibroblasts on Ru-LdECM hydrogels had an altered morphology as illustrated by the higher area (p < 0.0001, Mann-Whitney test) with an increased circularity (p < 0.0001, Mann-Whitney test) compared with the fibroblasts seeded on LdECM hydrogels (**Figure 8**).

**Figure 8:**
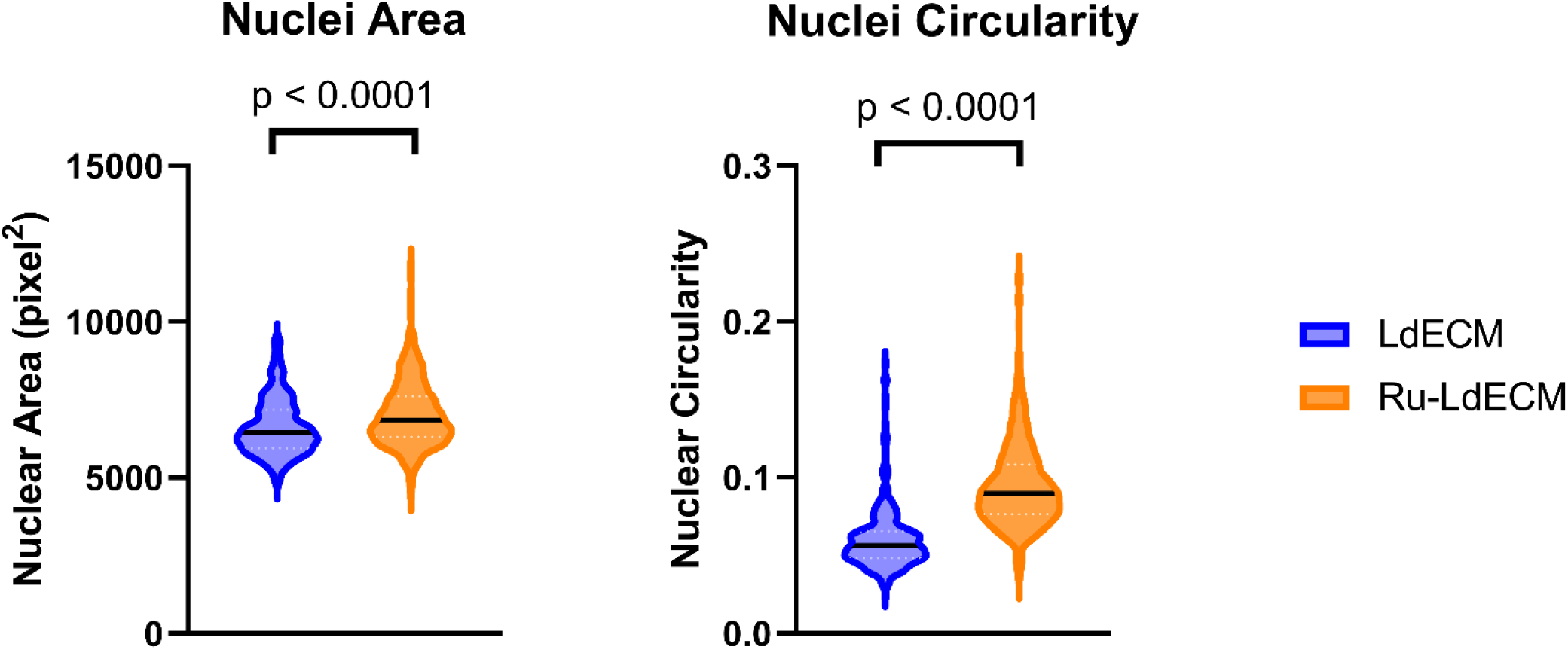
Comparison of the nuclear characteristics of fibroblasts seeded on LdECM and Ru-LdECM hydrogels. DAPI-stained fluorescent images of the fibroblasts were analysed using the CellProfiler software to compare the nuclear area and circularity. A) Comparison of nuclear area of fibroblasts seeded on LdECM and Ru-LdECM. B) Nuclear circularity of the fibroblasts seeded on LdECM and Ru-LdECM hydrogels. Each dot represents measurement on an individual nucleus for the respective characteristic, in total from 5 different randomized regions on the fluorescent images of DAPI staining for each sample (n=5). Applied statistical test: Mann-Whitney test. LdECM: Lung-derived ECM Hydrogels, Ru-LdECM: Ruthenium-crosslinked Lung-derived ECM Hydrogels

## Discussion

In this study we describe a model that enables modulation of the mechanical properties of an ECM without changing the composition. Using this model, we illustrated that by modulating the crosslinks between ECM fibres, the stiffness and stress relaxation properties of the ECM-derived hydrogels were altered. The crosslinking influenced the ECM fibre characteristics with a higher percentage of high density matrix and lower percentage of alignment being evident within the hydrogels treated with ruthenium. Fibroblasts grown on the surface of the crosslinked hydrogels displayed more myofibroblast-like characteristics. The features of this model illustrate that it would provide a novel research tool for investigating the importance of biomechanical changes in fibrotic diseases.

The increase in stiffness caused by ruthenium crosslinking in the LdECM hydrogels is similar to the increase in stiffness seen in fibrotic lung diseases such as idiopathic pulmonary fibrosis (IPF). ^33^ Booth et al. measured the stiffness of whole non-IPF and IPF human lungs before and after decellularization and found that the fibrotic regions of the IPF lung often reached a stiffness of 100 kPa or more, a vast increase compared to that of normal lung which has an average stiffness of 1.96 kPa. ^15^ The increased stiffness of IPF human lung was still present in hydrogels when compared to hydrogels generated from control human lungs, albeit proportionally reduced when compared to the intact lung tissue. ^14^ Recreating the (patho)physiological stiffness is essential to study the corresponding behaviour of cells during fibrotic diseases. ^34^ Ruthenium crosslinking on ECM-derived hydrogels provides an ideal opportunity to recreate the (patho)physiological stiffness. This is due to the fact that it does not require additional modifications on the ECM itself, enabling the modification of the mechanical properties while keeping the biochemical composition of the ECM constant. Thus these hydrogels mimic only the altered mechanical properties observed in fibrotic diseases. Ruthenium crosslinking relies on the crosslinking of the tyrosine amino acids and the potential for employing this strategy on the ECM-derived hydrogels has recently been demonstrated by Kim et al. ^25^ The ruthenium crosslinking of the ECM in our model allows us to recreate the (patho)physiological mechanical environment in a fibrotic lung. This method can most likely be adapted to generate tissue specific fibrotic environments that are representative of many organ microenvironments.

Ruthenium crosslinked LdECM hydrogels had a lower and more complex stress relaxation than uncrosslinked LdECM hydrogels, similar to that of IPF human lung compared to normal human lung. ^35^ The requirement of a fourth Maxwell element in modelling the stress relaxation of the Ru-LdECM hydrogels indicates a more complex stress relaxation, where most likely there is an extra contributing factor when compared to the uncrosslinked hydrogel. To date, attributing individual Maxwell elements to specific components of a hydrogel (such as water, small molecules, cells, or ECM) remains difficult in absence of a dedicated systematic study. But the fourth element Ru-LdECM hydrogels require to describe their relaxation profile would most likely be due to the secondary network formed through the ruthenium crosslinking. A similar difference in stress relaxation was found between control and fibrotic human lung ECM-derived hydrogels, showing three Maxwell elements in control hydrogels and 4 Maxwell elements in fibrotic hydrogels. ^29^ While the differences in species might complicate comparisons between porcine and human lung-derived hydrogels with respect to mechanical properties, the presence of a fourth Maxwell element in a Ru-LdECM hydrogels suggests that these hydrogels were able to resemble the stress relaxation behaviour of fibrotic lung ECM-derived hydrogels. Since in our study the base material is the same LdECM, the necessity of the fourth Maxwell element in modelling the stress relaxation is most likely due to the addition of ruthenium-induced crosslinks.

Ruthenium crosslinking of ECM hydrogels led to a more dense ECM network reflective of tissue changes in fibrotic diseases. Crosslinking the ECM fibres together in LdECM hydrogels resulted in lower fibre lengths and more dense matrix packed areas. In fibrotic lung diseases like IPF, the collagen network undergoes post-translational modifications and is crosslinked by the lysyl oxidase (LO) family of enzymes, and other enzymes, which leads to a more mature and organized collagen network. ^36, 37^ The denser matrix with a high degree of crosslinking is a key feature of fibrotic lung disease and protects the ECM from proteolysis. ^38^ The overall organization of the ECM in IPF is decreased when compared to normal lungs, a characteristic which was also present in the Ru-LdECM hydrogels as seen in the lower alignment. ^39^ Similar values in normalized numbers of endpoints and branchpoints suggest that fibre integrity was not affected during the crosslinking and existing branches were crosslinked. The shorter fibres in crosslinked hydrogels might be explained with the increased curvature in these samples: the effect of crosslinking on curvature of fibres with respect to the fibre length was prominent in shorter fibres (<40 μm) while longer fibres did not have differences among the two groups. These observations indicate that the ruthenium-crosslinking mainly influences the shorter fibres and decreases the average fibre length. Together with the mechanical characterization data, these results show that the mechanical properties were altered through the changes in the alignment and density of the matrix (HDM) in the crosslinked hydrogels.

One of the most important components of the fibrotic microenvironment is the (myo)fibroblasts and their responses to the altered ECM. In our model, the cells remained viable and the fibroblasts seeded on crosslinked hydrogels lost their spindle-like morphology that we observed on the native LdECM hydrogels. This observation is in parallel with previously reported studies showing the effect of stiffness on human lung fibroblasts. ^40^ As a response to the altered mechanical properties in Ru-LdECM hydrogels, the fibroblasts have displayed more myofibroblast-like characteristics, which is in reflective of previously published results. ^40^ The accompanying changes in the nuclear morphology was also in agreement with the influence of increased stiffness. ^41^ As reviewed by Wang et al., nuclear mechanotransduction as a response to a stiffer microenvironment (such as in fibrosis) has yet to be completely understood; however, such a change can result in altered gene regulation or nuclear transportation of cytoplasmic factors. ^42^ Another implication of the altered nuclear morphology due to increased stiffness was shown to influence the differentiation of mesenchymal stromal cells. ^43^ These demonstrated changes on the seeded fibroblasts in our study suggest that the biomechanical properties of the fibrotic microenvironment were replicated in our model.

Our study utilizes a crosslinking strategy on native ECM-hydrogels using a ruthenium complex and sodium persulfate, as described recently by Kim et al. ^25^ While this novel approach has its advantages, our study has also some limitations. In this study, we have seeded the fibroblasts on top of the hydrogels (2D) instead of seeding them within the hydrogel network. Although a 3D environment would represent the physiological situation in the body, a 2D culture system was preferred in this study to ensure proper visualization of the cell viability and morphology. Our study reports a model for examining the influence of biomechanical changes of the fibrotic microenvironment without investigating any gene and protein output from the fibroblasts. Although the influence of a fibrotic biomechanical microenvironment on fibroblasts have been shown to promote a pro-fibrotic phenotype both in gene and protein levels (as reviewed in ^5, 44^), such investigations are beyond the scope of this study. Lastly, the power of Maxwell modelling of the stress relaxation profiles of the native and crosslinked ECM hydrogels has not been completely realized and seems to remain as a mathematical exercise. The reason is that unlike other research areas e.g. microbial biofilms where relaxation constants has been linked to the composition ^45^, for hydrogels this systematic study is not yet available. With that, such modelling still proves useful in terms of analysing the altered stress relaxation behaviour.

## Conclusion

This study demonstrates the mechanical characterization of an *in vitro* ECM-based fibrosis model for advancement of investigations on effects of a fibrotic microenvironment on the cells. The next step for this model is to investigate how changes in the stiffness or viscoelastic relaxation can instruct the cells for further profibrotic responses, especially in a 3-dimensional setting. In addition, fibre characteristics analysis revealed that the changes in the fibre organization (alignment, density, curvature) accompany the altered pattern in the viscoelastic stress relaxation behaviour. More research on the influence of these altered fibre characteristics on the profibrotic activation of different cells (fibroblasts, macrophages…) have yet to be explored. Overall, this study shows the preparation and the characterization of an *in vitro* fibrosis model. Such advanced *in vitro* models for fibrosis research will improve our understanding on de-coupling the mechanical changes from the biochemical changes taking place in fibrosis.

## Supporting information

Electronic supplementary information

