## Supplementary material for "An *in vitro* model of fibrosis using crosslinked native extracellular matrix-derived hydrogels to modulate biomechanics without changing composition": Electronic supplementary information

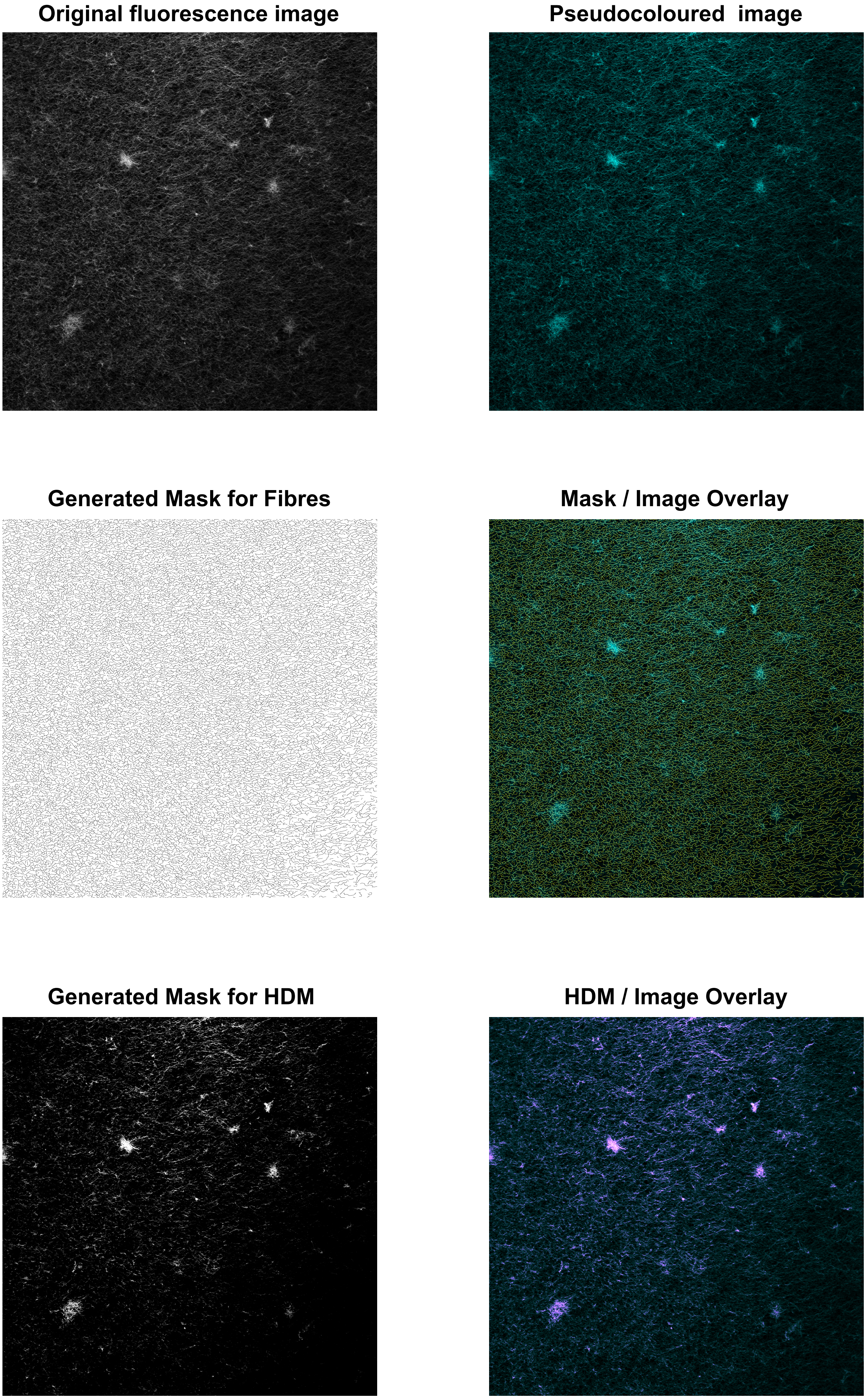


**Figure S1: Example of the masks and HDM generated through TWOMBLI analysis and their overlays with the original images**

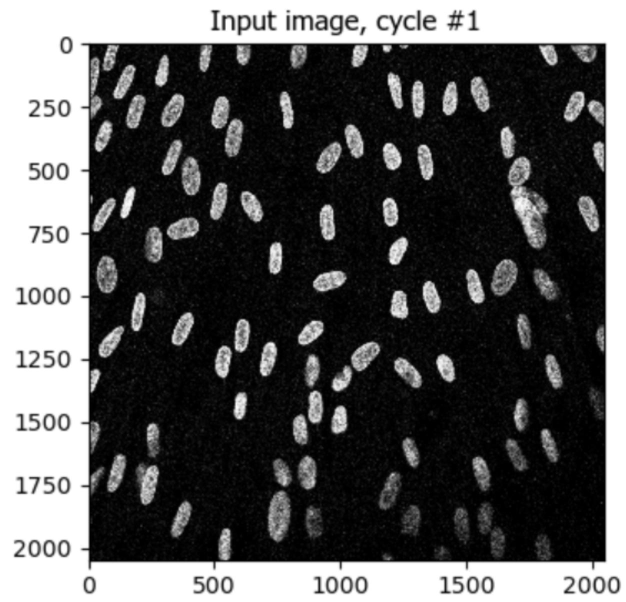


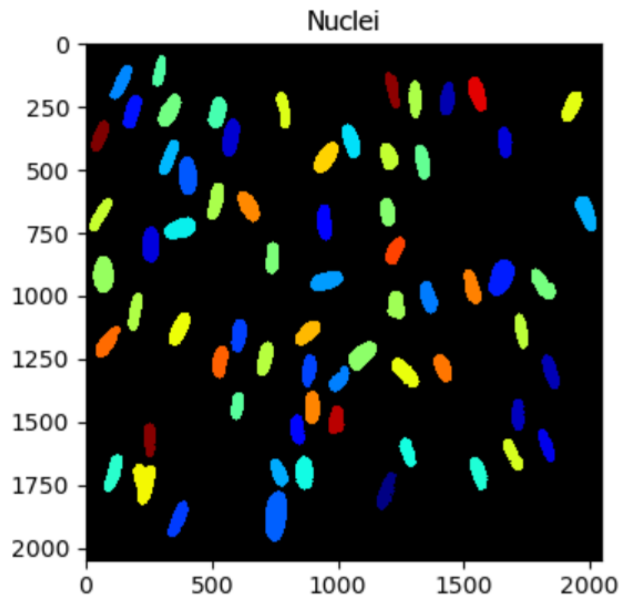


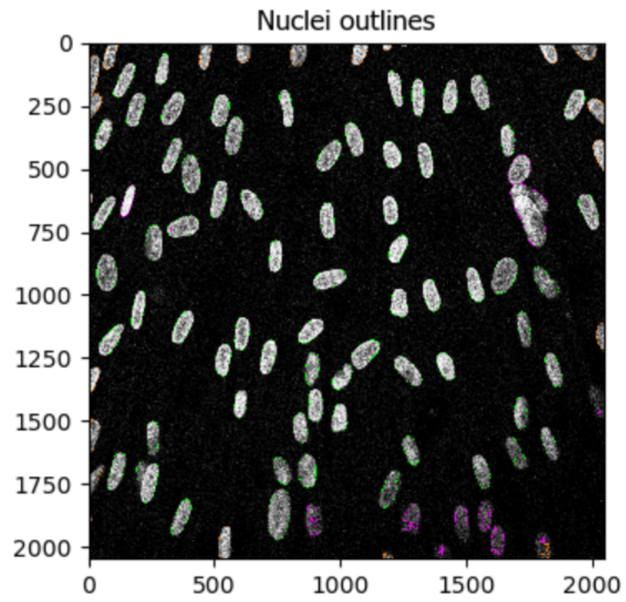


| | |
|---|---|
| # of accepted objects | 68 |
| 10th pctile diameter | 83.9 pixels |
| Median diameter | 91.7 pixels |
| 90th pctile diameter | 102.9 pixels |
| Area covered by objects | 11.1 % |
| Thresholding filter size | 1.0 |
| Threshold | 0.169 |
| Declumping smoothing filter size | 50.4 |
| Maxima suppression size | 52.5 |

**Figure S2: Example pipeline of the CellProfiler software for the nuclei analysis**

A)

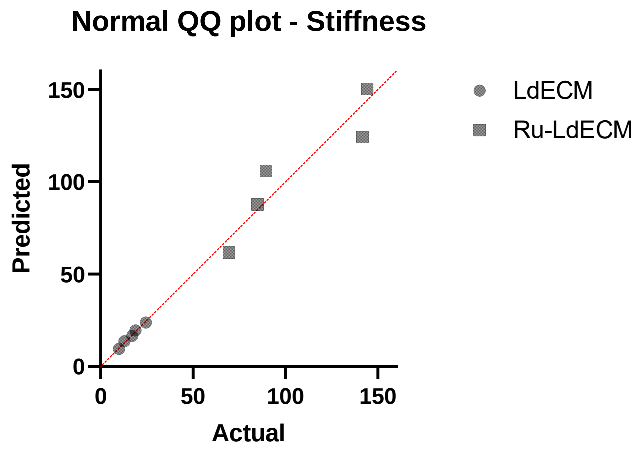
Normal QQ plot - Stiffness
Predicted
Actual
0
50
100
150
LdECM
Ru-LdECM

B)

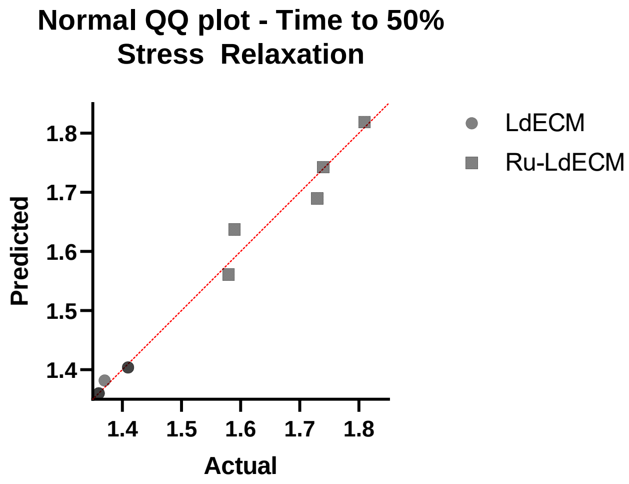
Normal QQ plot - Time to 50% Stress Relaxation
Predicted
Actual
1.4
1.5
1.6
1.7
1.8
LdECM
Ru-LdECM

C)

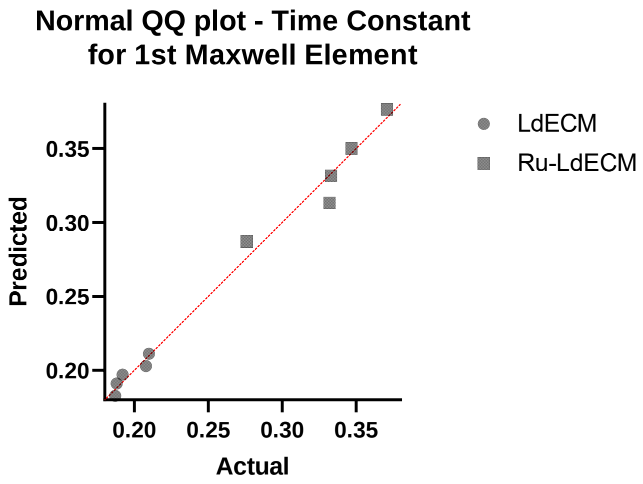
Normal QQ plot - Time Constant for 1st Maxwell Element
Predicted
Actual
0.20
0.25
0.30
0.35
LdECM
Ru-LdECM

D)

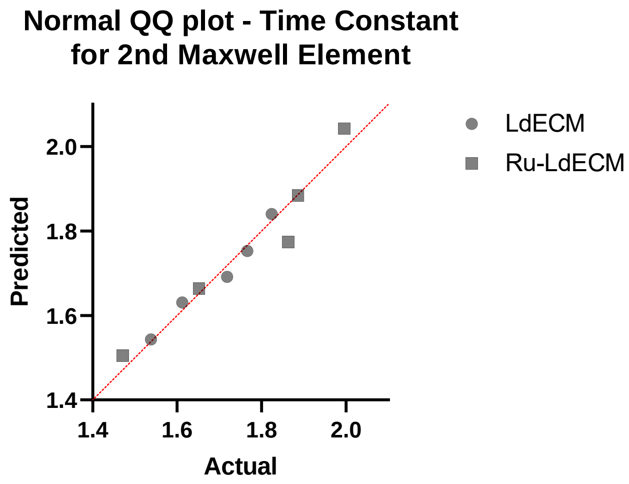
Normal QQ plot - Time Constant for 2nd Maxwell Element
Predicted
Actual
1.4
1.6
1.8
2.0
LdECM
Ru-LdECM

E)

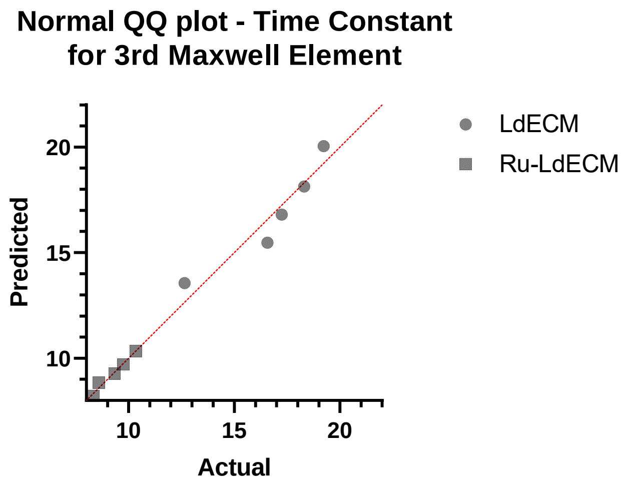
Normal QQ plot - Time Constant for 3rd Maxwell Element
Predicted
Actual
10
15
20
LdECM
Ru-LdECM

F)

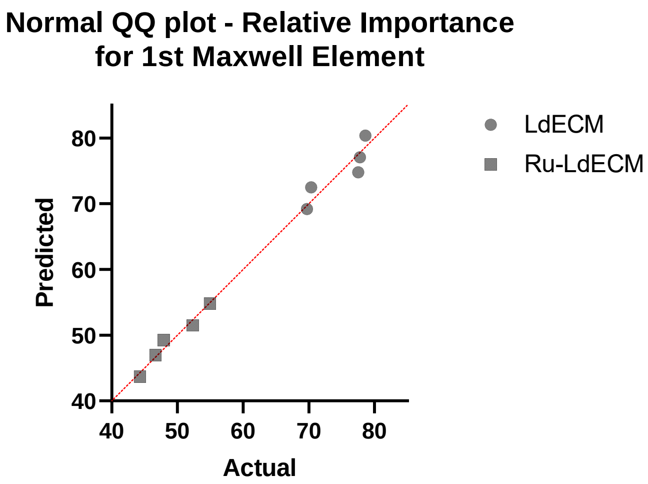
Normal QQ plot - Relative Importance for 1st Maxwell Element
Predicted
Actual
40
50
60
70
80
LdECM
Ru-LdECM

**G)**

**Normal QQ plot - Relative Importance for 2nd Maxwell Element**

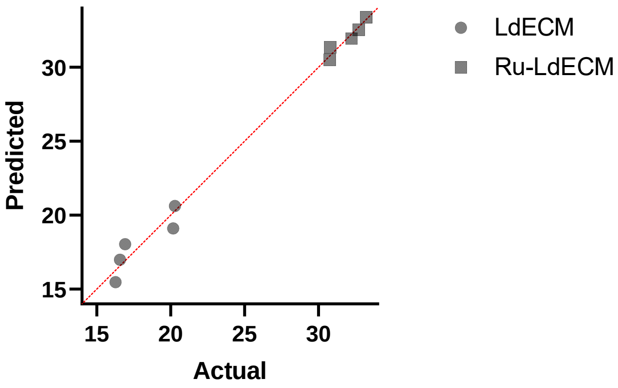


**H)**

**Normal QQ plot - Relative Importance for 3rd Maxwell Element**

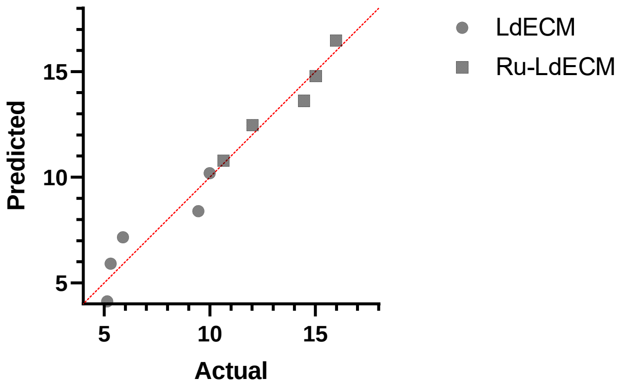


**I)**

**Normal QQ plot - Average Fibre Length**

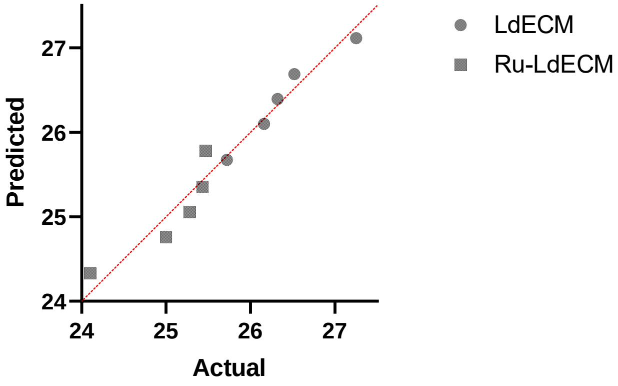


**J)**

**Normal QQ plot - Endpoints**

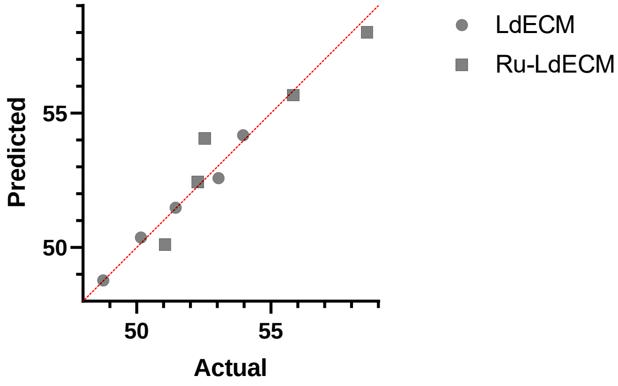


**K)**

**Normal QQ plot - Branchpoints**

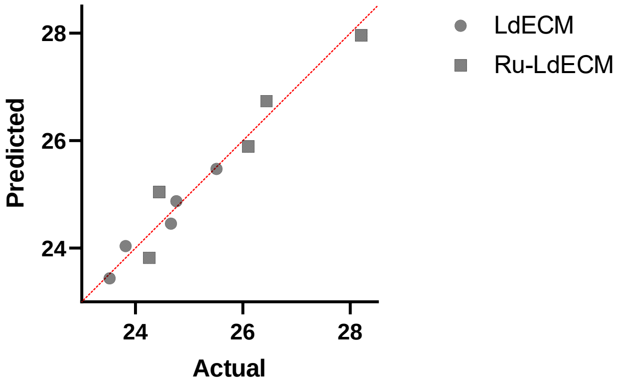


**L)**

**Normal QQ plot - Hyphal Growth Unit**

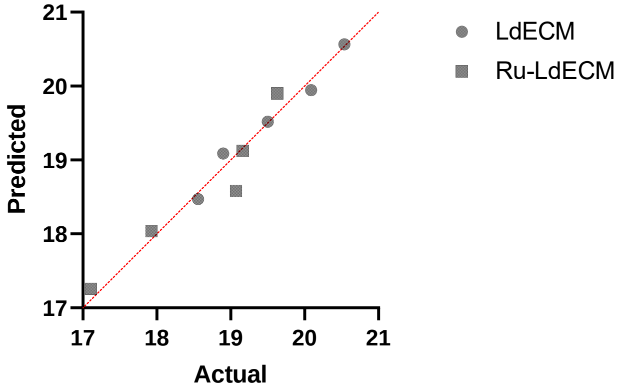

**M)**

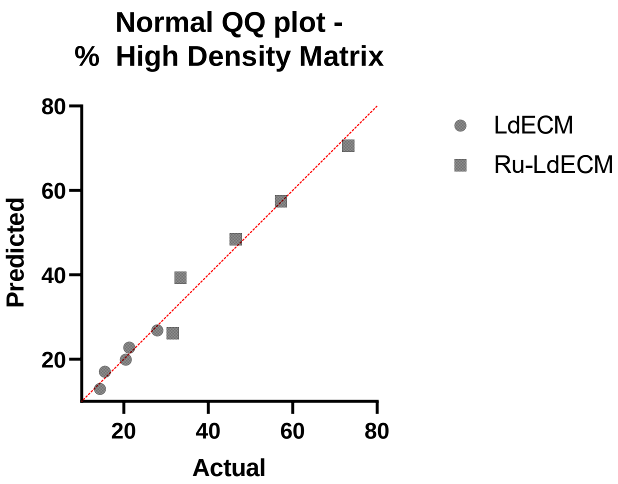
Normal QQ plot -
% High Density Matrix
Predicted
80
60
40
20
20
40
60
80
Actual
LdECM
Ru-LdECM

**N)**

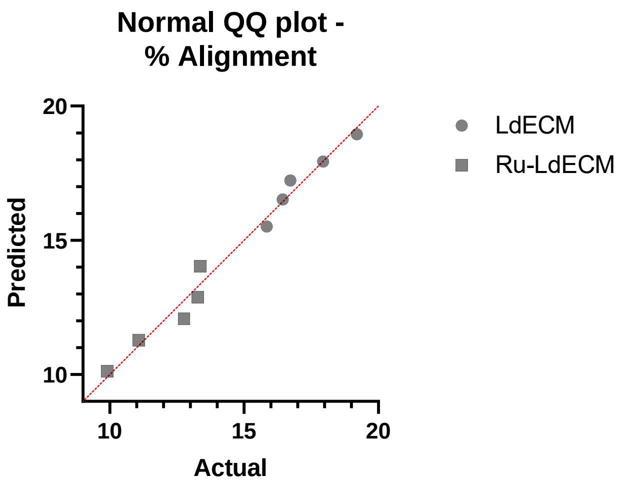
Normal QQ plot -
% Alignment
Predicted
20
15
10
10
15
20
Actual
LdECM
Ru-LdECM

**O)**

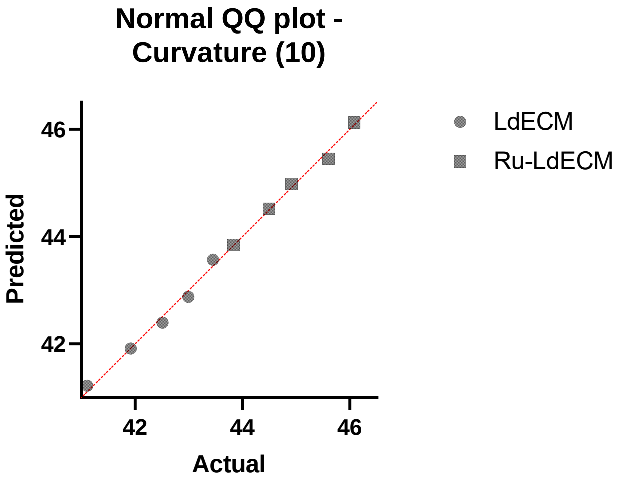
Normal QQ plot -
Curvature (10)
Predicted
46
44
42
42
44
46
Actual
LdECM
Ru-LdECM

**P)**

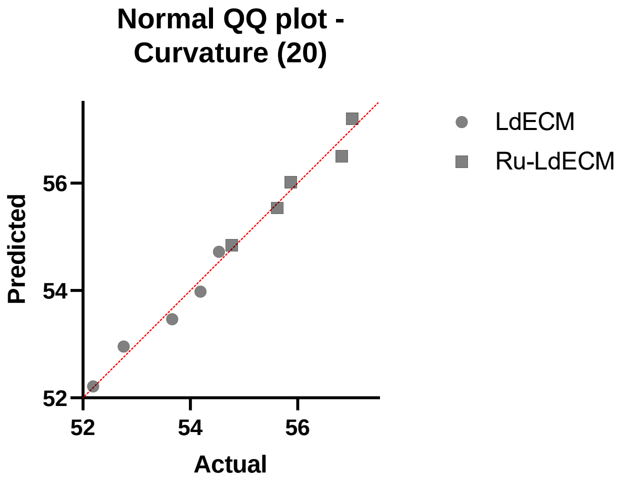
Normal QQ plot -
Curvature (20)
Predicted
56
54
52
52
54
56
Actual
LdECM
Ru-LdECM

**R)**

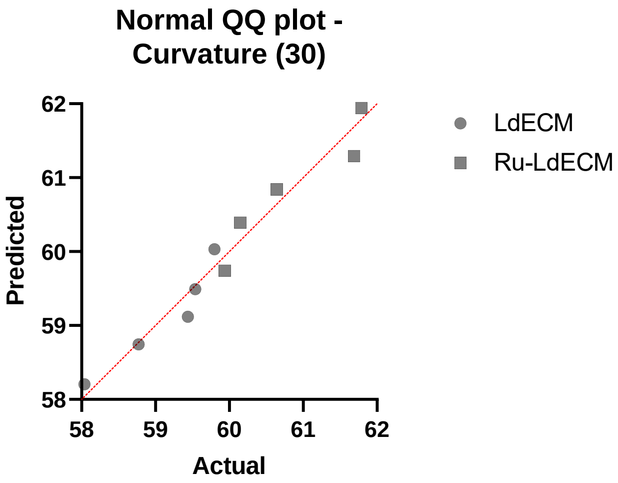
Normal QQ plot -
Curvature (30)
Predicted
62
61
60
59
58
58
59
60
61
62
Actual
LdECM
Ru-LdECM

**S)**

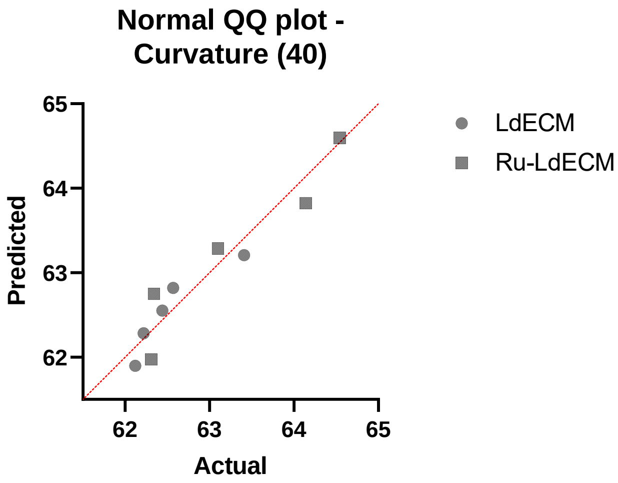
Normal QQ plot -
Curvature (40)
Predicted
65
64
63
62
62
63
64
65
Actual
LdECM
Ru-LdECM

**T)**

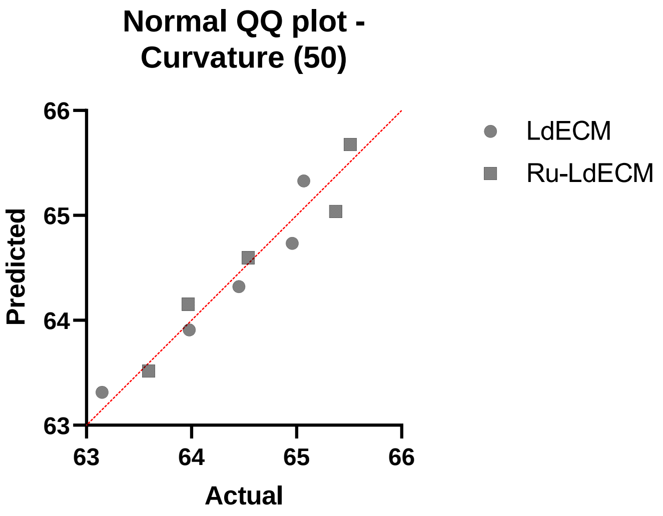


**Figure S3: Q-Q plots for the normality assessments of the data analyzed using parametric tests. A) Stiffness, B) Time to 50% Stress Relaxation, C) Time Constant for 1st Maxwell Element, D) Time Constant for 2nd Maxwell Element, E) Time Constant for 3rd Maxwell Element, F) Relative Importance for 1st Maxwell Element, G) Relative Importance for 2nd Maxwell Element, H) Relative Importance for 3rd Maxwell Element, I) Average Fibre Length, J) Number of Endpoints, K) Number of Branchpoints, L) Hyphal Growth Unit, M) % High Density Matrix, N) % Alignment, O) Curvature (10 μm fibres), P) Curvature (20 μm fibres), R) Curvature (30 μm fibres), S) Curvature (40 μm fibres), T) Curvature (50 μm fibres).**

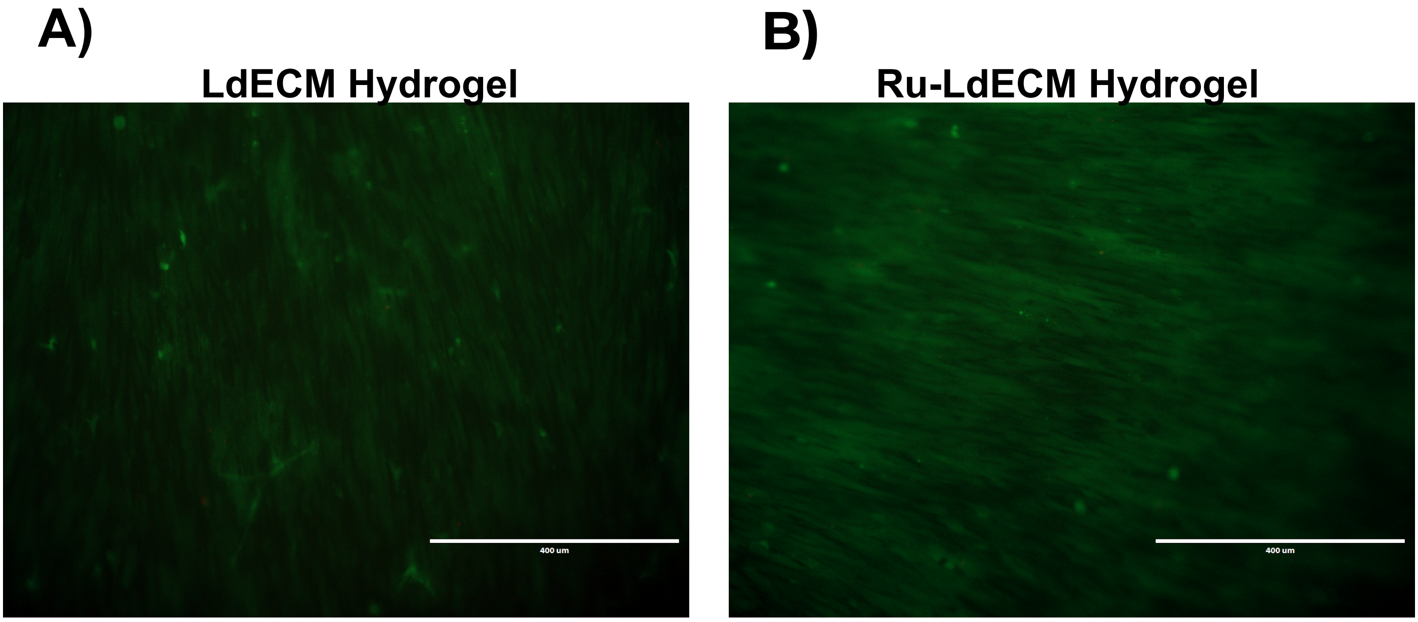


**Figure S4: Example image of the live/dead staining at day 7**
